# Anti-correlation of LacI association and dissociation rates observed in living cells

**DOI:** 10.1101/2024.08.21.608596

**Authors:** Vinodh Kandavalli, Spartak Zikrin, Johan Elf, Daniel Jones

**Affiliations:** Department of Cell and Molecular Biology, Science for Life Laboratory, Uppsala University, Husargatan 3, Uppsala 75124, Sweden

**Keywords:** Transcription factor dynamics, association and dissociation rates, Cell-to-cell variability, mRNA FISH, DNA-protein interaction, Gene expression noise

## Abstract

The rate at which transcription factors (TFs) bind their cognate sites has long been assumed to be limited by diffusion, and thus independent of binding site sequence. Here, we systematically test this assumption using cell-to-cell variability in gene expression as a window into the *in vivo* association and dissociation kinetics of the model transcription factor LacI. Using a stochastic model of the relationship between gene expression variability and binding kinetics, we performed single-cell gene expression measurements to infer association and dissociation rates for a set of 35 different LacI binding sites. We found that both association and dissociation rates differed significantly between binding sites, and moreover observed a clear anticorrelation between these rates across varying binding site strengths. These results contradict the long-standing hypothesis that TF binding site strength is primarily dictated by the dissociation rate, but may confer the evolutionary advantage that TFs do not get stuck in near-operator sequences while searching.

## Introduction

Cell-to-cell variability in gene expression, sometimes called “noise,” is a fact of life in the low molecular copy number regime in bacterial cells. A variety of mechanisms underpin this variability, including the stochastic binding and unbinding of transcription factors^1^ and RNA polymerase (RNAP)^2^, gene dosage effects, and partitioning of macromolecules upon cell division^3,4^, among many others^5,6^. Conversely, measurements of gene expression variability can be used to shed light on these cellular processes^7–9^, although attributing observed variability to specific mechanisms can be challenging. Among the biological implications of gene expression variability, one of the most prominent is “bet-hedging,” the idea that subpopulations of cells exhibiting altered gene expression states may be better prepared to survive rapid and unpredictable shifts in environmental conditions^10,11^.

In this work, we focus on variability due to the stochastic association and dissociation of transcription factors (TFs). A longstanding assumption has been that TF association is diffusion-limited: that is, that the association rate is largely independent of the strength of the transcription factor binding site^7,13^. Under this assumption, the strength (reflected in the dissociation constant *K*_*d*_) of a particular binding site is modulated by changes in the dissociation rate *k*_*d*_. However, recent *in vitro* experiments have called this assumption into question, showing that the dissociation rate *k*_*d*_ and association rate *k*_*a*_ were negatively correlated for the model transcription factor LacI^14^.

Here, we use cell-to-cell variability in gene expression as a window into LacI kinetics in living cells. By modeling the relationship between variability and LacI kinetics, we use single-cell gene expression measurements to infer association and dissociation rates for a set of 35 different LacI binding sites. Consistent with recent *in vitro* results, we find a clear anticorrelation between association and dissociation rates, upending longstanding understandings of *in vivo* transcription factor binding kinetics.

## Results

We designed a set of promoter-operator constructs in which LacI binding sites were placed immediately downstream of the apFAB120 promoter (Supplementary Fig. 1), such that transcription from the promoter is occluded when LacI is bound, and transcription is enabled when LacI is not bound (Fig. 1A). To relate the observed mRNA copy number distributions to LacI association and dissociation rates, we used the standard two-state model of stochastic gene expression, sometimes referred to as the “random telegraph” model ^12,15^ (Fig. 1A). In this model, the promoter can be found in one of two states depending on whether LacI is bound to its operator. In the “on” state, corresponding to unbound LacI, transcripts are produced at rate *r*; in the “off” state corresponding to bound LacI, transcription is blocked. In both states, mRNAs are degraded at rate *γ*. Transitions from “off” to “on” occur at rate *k*_*d*_, corresponding to LacI dissociation, whereas transitions from “on” to “off” occur at rate *k*_*a*_, corresponding to LacI association.

**Fig. 1:**
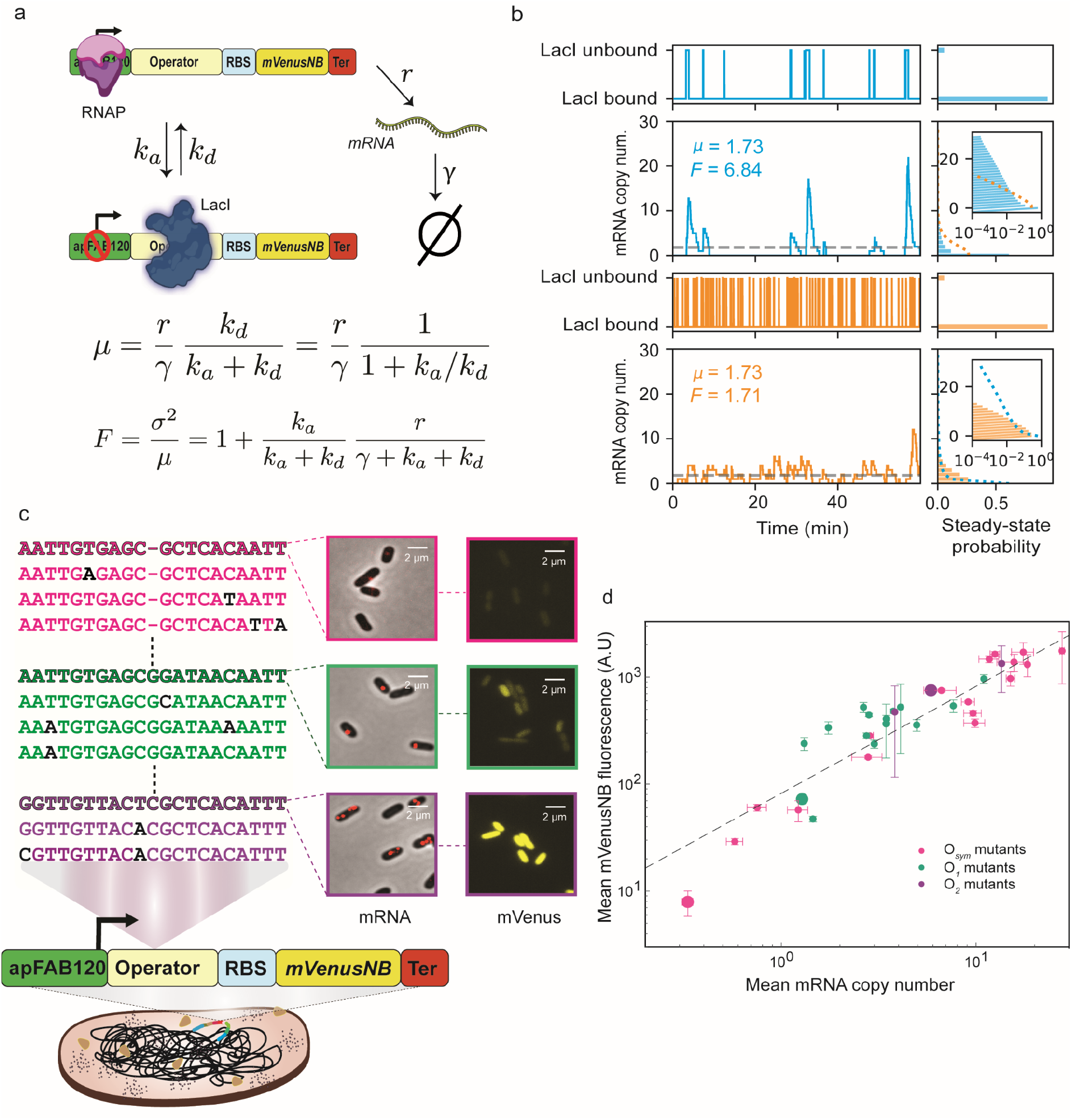
Promoter-operator construct and model relating gene expression variability to transcription factor kinetics. **a** The construct consists of a LacI binding site (“Operator”) immediately downstream of the apFAB120 promoter, driving expression of the mVenusNB fluorescent protein. The system is either in a LacI-unbound state (top) in which mRNA is produced at rate *r*, or a LacI-bound state in which transcription is halted (bottom). mRNA is degraded at rate *γ*, and *k*_*a*_ and *k*_*d*_ represent the LacI association and dissociation rates, respectively. Expressions for mean expression μ and Fano factor F as a function of model parameters are shown below the schematic^12^. **b** Stochastic simulations of mRNA production for “slow” (top, blue; *k*_*a*_=3.0 min^-1^, *k*_*d*_=0.18 min^-1^) and “fast” (bottom, orange; *k*_*a*_=30 min^-1^, *k*_*d*_=1.8 min^-1^) operators. *r*=25 min^-1^ and *γ*=0.8 min^-1^ for both operators. Both operators have the same steady-state LacI binding probability and hence the same mean mRNA expression. However, the “slow” operator exhibits greater variability, reflected in its larger Fano factor and longer-tailed steady-state mRNA copy number distribution. For easier comparison, the “fast” mRNA distribution is plotted (purple dashed line) on top of the “slow” mRNA distribution (green bars) and vice versa. The mRNA distributions are plotted on a log scale in insets. **c** Subset of operator mutants assayed in this paper. Operators used the O_*sym*_ (pink), O_*1*_ (green), or O_*2*_ (purple) operators as starting points. The wild-type operators are shown in bold and mutations are highlighted in black. Gene expression levels as measured by mRNA FISH (“mRNA”) and fluorescence microscopy (“mVenusNB”) are shown for O_sym_, O_*1*_, and O_2_. **d** For each operator, the mean mRNA copy number is plotted against the mean mVenusNB fluorescence, showing (as expected) a strong correlation. O_*sym*_, O_1_, and O_2_ wild-type operators are plotted with larger circles; error bars represent sem from bootstrapping.

A full analytic solution to the resulting chemical master equation has previously been derived^12,15^. Under this model, the mean mRNA expression is given by

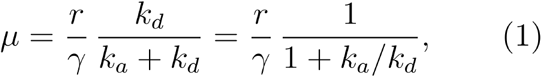

where 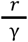 is the mean mRNA copy number in the absence of LacI while the second term is simply the fraction of time that LacI is not bound. Notably, the mean expression is dependent on the ratio of association rate to dissociation rate, but does not depend on their absolute magnitude.

To obtain the absolute magnitudes of *k*_*a*_ and *k*_*d*_, we need to consider higher moments of the distribution. It is convenient to use the Fano factor, defined as the variance *σ*^2^ divided by the mean μ. For this model, the Fano factor is given by

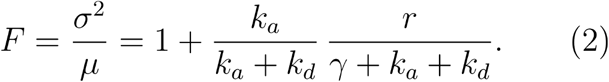

In Fig. 1B, stochastic simulations of two hypothetical operators are shown: one operator with slow association and dissociation kinetics (top, blue), and one operator with fast kinetics (bottom, orange), as well as the corresponding steady-state distributions (Fig. 1B, right). Both operators have the same *k*_*a*_ to *k*_*d*_ ratio, resulting in the same fractions of time spent in the “on” (LacI unbound) and “off” (LacI bound) states and the same mean expression. However, the slow operator is characterized by infrequent bursts of large numbers of transcripts, whereas the fast operator maintains a more constant production. This difference is reflected in a broader steady-state distribution and a larger Fano factor value for the slow operator.

To systematically investigate the relationship between association and dissociation rates for LacI binding sites, we selected a set of 35 binding sites to span a broad range of binding site strengths (Fig. 1C)^14^. We used the synthetic O_sym_ operator as well as the naturally occurring O_1_ and O_2_ operators as starting points, and constructed mutants of each operator (Supplementary table 1). These promoter-operator constructs were used to drive expression of the fluorescent protein mVenusNB (Fig. 1C).

For each operator, we quantified gene expression using single-cell mRNA FISH (Fig. 1C, “mRNA”, Supplementary Fig. 2), allowing us to determine the distribution of mRNA copy numbers across a population of fixed cells. As expected, the wide range of binding site strengths was reflected in mean mRNA copy number values ranging from 0.3 to 20 (Supplementary Fig. 3). In parallel, we measured mVenusNB fluorescence for each construct (Fig. 1C, “mVenusNB”) and found that mean mRNA and mean protein fluorescence were highly correlated across our panel of LacI binding sites, as expected (Fig. 1D). We also validated the mRNA FISH data with relative RT-qPCR measurements for a subset of binding sites, and found excellent agreement between the measurements (Supplementary Fig. 4).

### Calculation of *k*_*a*_ and *k*_*d*_ from mRNA statistics

In Fig. 2, the procedure for calculation of *k*_*a*_ and *k*_*d*_ is illustrated using the O_*sym*_ operator as an example. Equations (1) and (2) contain two unknowns besides *k*_*a*_ and *k*_*d*_: the degradation rate γ and the basal transcription rate *r*. The degradation rate γ was obtained for the O_sym_, O_1_, and O_2_ operators by halting transcription at t=0 using rifampicin (500 μg/mL) and measuring mRNA levels at subsequent time points using bulk RT-qPCR, yielding a value of γ = 0.79 min^-1^ (Fig. 2A, left), consistent with previously published results ^16,17^. The basal transcription rate *r* was determined separately for each operator by measuring the mean mRNA copy number in Δ*lacI* strains and using the fact that *μ*_Δ*lacI*_ = *r / γ*, enabling estimation of r once the degradation rate γ and mean expression *μ*_*ΔlacI*_ are known (Fig. 2A, right). For the O_*sym*_ operator, we estimated that *r* = 24.6 min^-1^. With *r* and *γ* known, and the mean and Fano factor experimentally determined using mRNA FISH, Equations (1) and (2) constitute a system of two equations with two unknowns and can be numerically solved for *k*_*a*_ and *k*_*d*_. Before doing so, we correct the Fano factor for the effect of RNAP copy number variability to obtain the corrected Fano factor *F*_*c*_ = *F* - *μ* / 10, as described previously^18,19^, then use *F*_*c*_ in place of F when solving for k_a_ and k_d_ (Fig. 2b). The inferred k_a_ and k_d_ values are substantively unchanged even if this correction is not included (Supplementary Fig. 5). For both Δ*lacI* and *wt lacI* experiments, only cells with lengths between 1.73 and 2.16 μm were included in the analysis (out of a length range of approximately 1.4 to 3.25 μm across all cells), to ensure that only cells with a single copy of the promoter-operator construct were present and thus counteract variability from gene copy number variation^18,19^.

**Fig 2:**
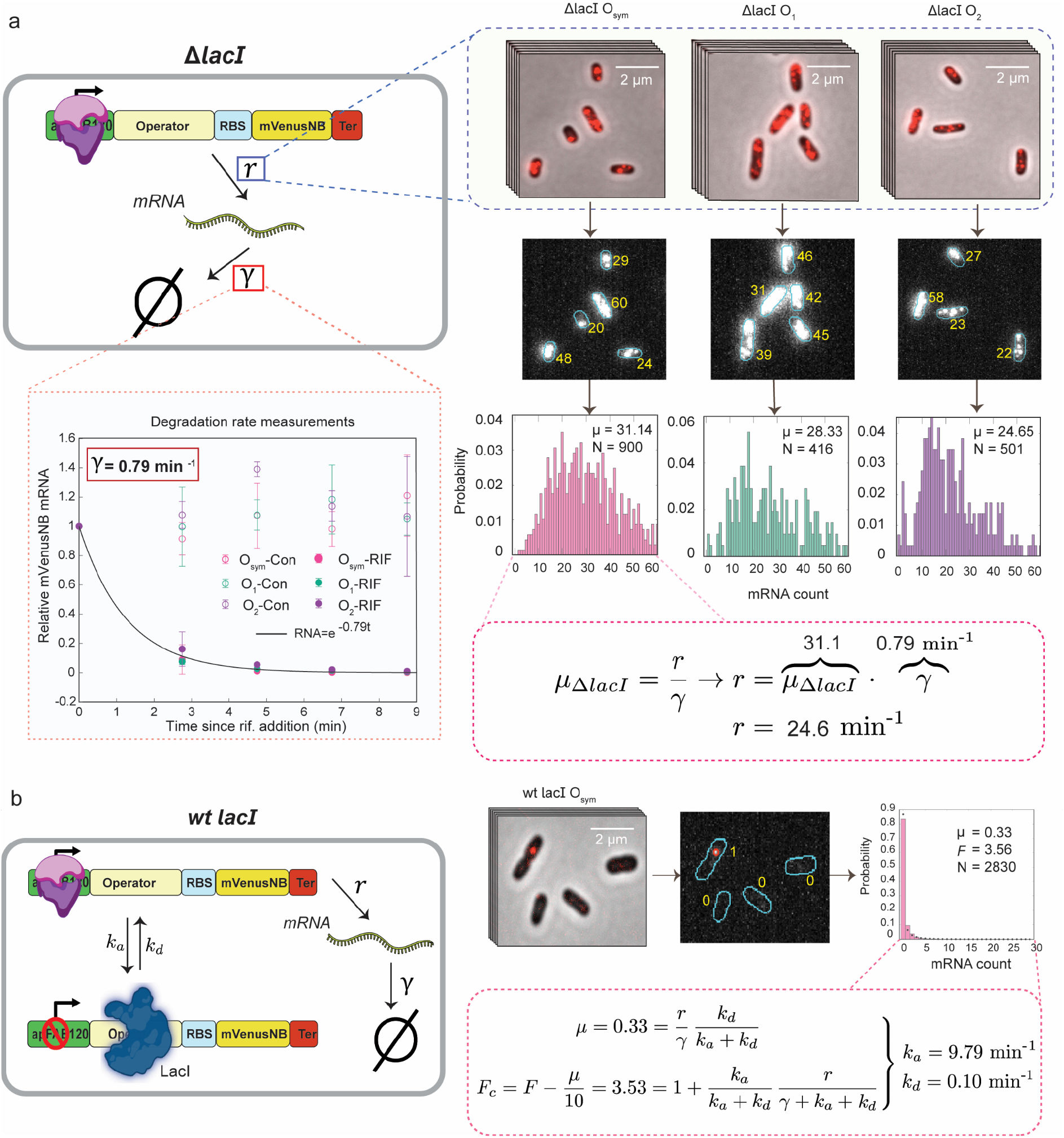
Calculation of *k*_*a*_ and *k*_*d*_ for O_sym_ operator. **a** First, *r* and *γ* are determined in a strain in which LacI is deleted. γ is determined by performing bulk RT-qPCR on the mVenusNB mRNA (3 biological replicates) on samples taken at two-minute intervals following addition of rifampicin to halt transcription. Data points labeled “RIF” correspond to rifampicin-treated samples while points labeled “con” represent untreated samples. *r* is determined by measuring mean mRNA expression *μ*_*ΔlacI*_ in Δ*lacI* strains using mRNA FISH, and using the equation *μ*_*ΔlacI*_ *= r / γ*. **b** Once *r* and *γ* are known, *k*_*a*_ and *k*_*d*_ are determined by measuring the mRNA copy number distribution in cells expressing LacI. From the distribution, the mean μ and Fano factor *F* are computed. The Fano factor is corrected for the effect of RNAP copy number variation (see main text) to obtain the corrected Fano factor *F*_*c*_, whereupon the appropriate values for μ, *F*_*c*_, *r*, and *γ* are substituted in Eq. 1 and 2, which are solved numerically for *k*_*a*_ and *k*_*d*_.

This procedure was repeated to estimate *k*_*a*_ and *k*_*d*_ for each of the 35 LacI binding sites. The value of *r* was estimated individually for each operator from Δ*lacI* experiments, in order to account for potential differences in basal transcription rates due to changes in operator sequence, which in turn affects the 5’ UTR and hence potentially transcription initiation. We found that changes in operator sequence weakly affected transcription in Δ*lacI* strains, with a maximum difference of about twofold (Supplementary Fig. 6). In Fig. 3a, the experimentally-determined mRNA copy number distributions are shown for the O_sym_, O_1_, and O_2_ operators (colored histograms), along with corresponding model predictions given the estimated values of *k*_*a*_ and *k*_*d*_ (black dotted lines), which show good agreement. The corresponding distributions are shown in Supplementary Fig. 3 for all operators. In Fig. 3b, the estimated *k*_*a*_ values are plotted against *k*_*d*_ for all operators, revealing a distinct anticorrelation between these two rates, and demonstrating that changes in operator strength are reflected in both rates, rather than primarily in *k*_*d*_. In Supplementary Fig. 7, we compare our results with previous *in vitro* measurements of the same operators^4^ and find reasonable agreement between *in vitro* operator occupancy and *in vivo* repression ratio (Supplementary Fig. 7a), as well as between *in vitro* and *in vivo* rate measurements (Supplementary Fig. 7b, 7c).

**Fig 3:**
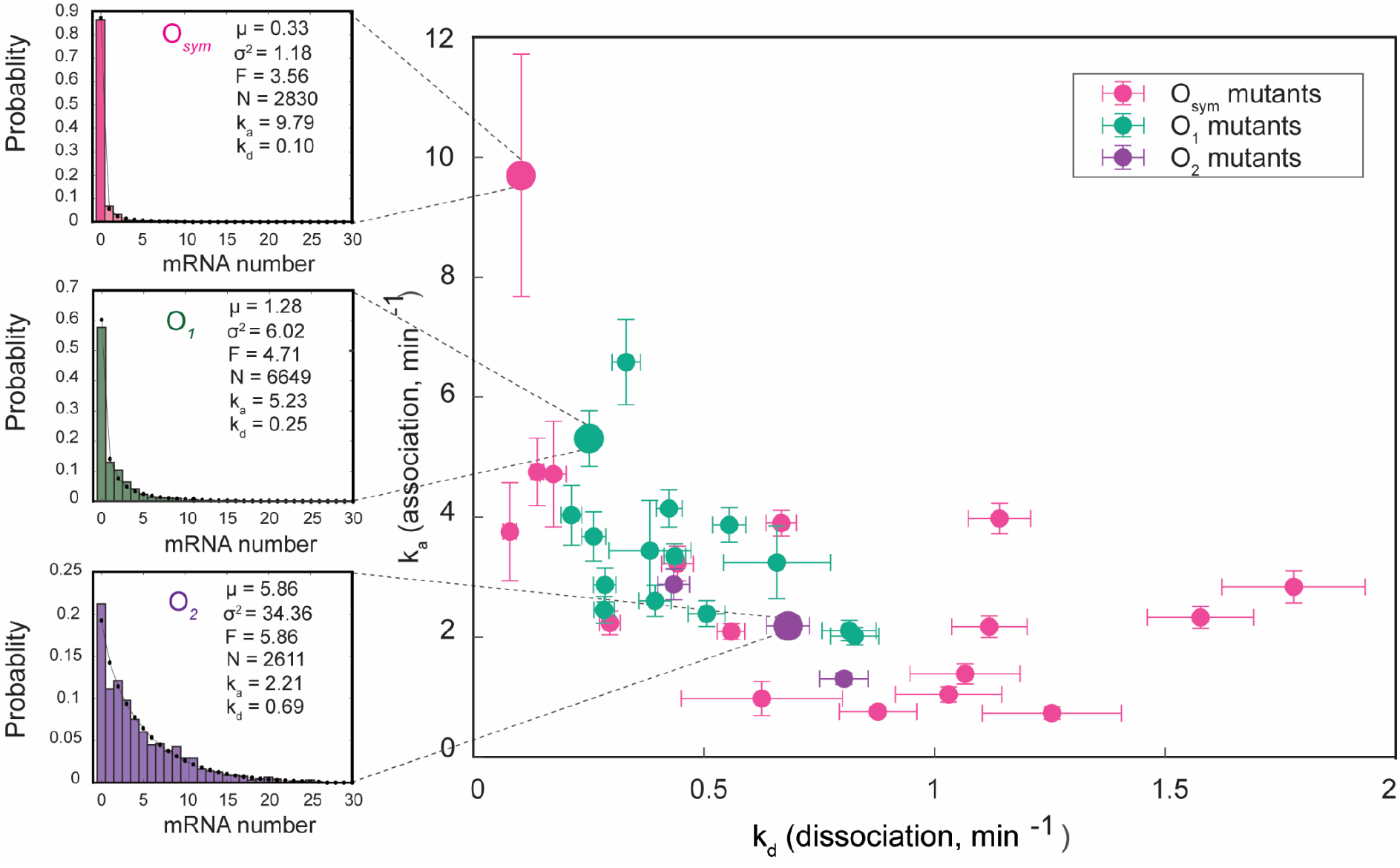
Selected mRNA distributions and inferred rates for all operators. **a** Experimentally observed mRNA copy number distributions are shown for the O_*sym*_, O_*1*_, and O_*2*_ operators, along with the model prediction given inferred rate parameters (black dots). **b** Association rate plotted against dissociation rate for all 35 operators assayed, showing a pronounced anticorrelation. Error bars represent SEM obtained by bootstrapping.

## Discussion

Here, we combined a mathematical model of stochastic gene expression with measurements of steady-state mRNA copy number distributions, to infer the LacI association and dissociation rates for a wide range of operator strengths. This system can be described with more complex models explicitly incorporating *e*.*g*. RNAP binding kinetics and open complex formation^7,20–22^, but our aim was to employ the simplest model that adequately described the observed distributions. For the *O*_*sym*_ operator, we obtained a dissociation rate of *k*_*d*_ = 0.10 min^-1^, in excellent agreement with *k*_*d*_ = 0.11 min^-1^ as obtained in a previous *in vivo* study using an alternative method^23^, strengthening the credibility of our approach. Our measurements also show reasonable agreement with previous *in vitro* results^14^ (Supplementary Fig. 7). Interestingly, *O*_*sym*_ variants appear to exhibit relatively lower *k*_*a*_ values and higher *k*_*d*_ values *in vivo* than *O*_*1*_ and *O*_*2*_ variants with similar *in vitro* rates (Supplementary Fig. 7). It is tempting to relate this discrepancy to the fact that *O*_*sym*_ is a synthetic operator and thus performs worse *in vivo* for reasons yet to be uncovered.

To our knowledge, this paper provides the first direct *in vivo* evidence of an anticorrelation between TF association and dissociation rates. Such anticorrelation appears if binding strength mainly is governed by the probability of recognizing and binding upon reaching the operator, which also shows up in the macroscopic rate of leaving the operator *k*_*d*_ since many rebinding events occur for strong operators^14^. This picture is contrary to the longstanding hypotheses that TF binding site strength is primarily dictated by the dissociation rate, but has the evolutionary advantage that TFs do not get stuck in near-operator sequences while searching.

At the same time, finding an appropriate regime in parameter space for basal transcription rate *r*, repressor concentration, and operator strength in order to carry out these measurements was not trivial. For strong operators such as O_sym_, *r* needed to be large enough compared to the association rate *k*_*a*_ for multiple transcripts to have a reasonable chance of being produced during LacI unbinding events. Otherwise, if each LacI unbinding event leads to 0 or 1 transcripts, transcription events become uncorrelated and hence effectively Poissonian, with the Fano factor equal to one. In this scenario, the variability carries no information about LacI kinetics. Conversely, too large an *r* value risks perturbing LacI binding kinetics as RNAP clears the promoter region, potentially hindering LacI from binding^21^. In Supplementary Fig. 8, we investigate this effect and find that it does not materially affect the conclusions of this paper. It was also necessary to balance LacI concentration with operator strength such that operators are neither always bound nor never bound, situations in which gene expression variability again carries little information about LacI kinetics (Supplementary Fig. 9). As the operators investigated here roughly spanned the range of operator strengths found in nature (from stronger than O_1_ to weaker than O_2_), the wild-type LacI concentration (*ca* 5-10 tetramers per cell^24^) proved to be an appropriate choice.

Nonetheless, the implication that operator strength is governed primarily by the recognition probability depends only on the observation of an anticorrelation between *k*_*a*_ and *k*_*d*_, regardless of the specific parameter regime required to observe this anticorrelation using our gene-expression-variability-based approach. Moreover, despite fine-tuning, none of the biological parameters in this study fall outside of the range observed in nature. The promoter architecture studied here (“simple repression”) is likewise one of the most common in *E. coli*^25^ and these results shed light on how cell-to-cell variability is affected by changes in binding site sequence for this ubiquitous motif. A complete understanding of the interplay between TF binding site strength, TF kinetics, gene expression variability, and the evolution of transcriptional regulatory sequences remains a compelling challenge for ongoing research.

## Methods

### Strain construction

*Chromosomal knockout of lacI gene*: The *lacI* gene was deleted using the DIRex method^26^. PCR fragments were amplified from the Acatsac1 (GenBank: MF124798) and Acatsac3 cassettes (GenBank: MF124799) using the oligos Del_LacI_Direx-F cat_mid-R, cat_mid-F, and Del_LacI_Direx-R (see supplementary table 2 for sequences). The PCR products were purified and electroporated into EL330 [*E. coli* K-12 MG1655] cells expressing lambda red proteins from plasmid pSIM5-Tet. After electroporation, cells were incubated in 1 mL of SOC medium at 30 °C for 1 h and then plated onto LB + 25 μg/mL chloramphenicol plates. The colonies were re-streaked on LB + 5% sucrose plates to select for cells with the desired recombination. The resulting colonies were confirmed by PCR. Further sequence confirmation was done for the deletion of LacI gene in the chromosome and hereafter the strain is referred to as EL4110 [*E. coli* K-12 MG1655, *ΔlacI*].

### Operator mutation library construction

Using Golden Gate cloning, we assembled a plasmid with an R6K origin of replication backbone and a Biopart that consists of the fluorescent protein (mVenusNB) with a strong ribosome binding site (RBS) downstream of the *lac* operator site, with expression driven by the constitutive synthetic promoter apFAB120^27^. Upstream of the promoter sequence, an ampicillin resistance cassette was introduced transcribing in the opposite direction (Supplementary Fig. 1).

Next, we amplified the Biopart with *ygaY* homology arms using the oligos ygaY_HA_F, and ygaY_HA_R, in preparation for chromosomal integration into the *ygaY* locus. The PCR fragment was further purified and electroporated into the EL330 and EL4110 strains producing lambda red proteins from plasmid pSIM5-tet. The successful chromosomal integration of the Biopart in the *ygaY* locus was confirmed by PCR and sequencing. For the operator mutation library, we selected single- and double-point mutations in *O*_*sym*_, *O*_*1*_ *and O*_*2*_ from mutants characterized in Marklund *et al*^14^ . Using the DIRex method, we constructed a total of 35 operator variants (see supplementary table 1) in both the EL330 and EL4110 strains. All these strains were also confirmed by sequencing.

### Growth medium and conditions

From the glycerol stocks (ࢤ80°C), cells were streaked on the fresh LB plates and grown overnight at 37 °C. Single colonies were inoculated in a LB medium and grown overnight in a continuous orbital shaking incubator with 200 r.p.m at 30 °C. From the overnight cultures, cells were further diluted in 1: 100 times in fresh M9 medium (containing M9 salts, 2 mM MgSO4, 0.1mM CaCl2, 0.4% succinate as a carbon source, supplemented with 0.5X RPMI). Cells were grown at 37 °C with 200 r.p.m until they reached mid-exponential phase (OD_600_ = ∼0.4). Next, cells were collected for microscopy experiments to quantify mRNA and protein expression levels.

### FISH probe design

To detect the mRNA of *mVenusNB* gene, 30 fluorescent DNA probes were designed using the Stellaris® Probe Designer version 4.2 (https://www.biosearchtech.com/stellaris-designer) and purchased from Integrated DNA Technologies (IDT Iowa, USA). Each DNA probe was 20 nt in length with a minimum distance of 2 nt and the masking level at 1–2. All probes were labeled by Cy-5 at the 3’ end of the DNA. Probe sequences are listed in supplementary table 3.

### FISH protocol

We followed the FISH protocol as in refs.^16,28^. Briefly, when cell cultures reached mid-exponential phase (OD_600_ ≅ 0.4), cells were collected by centrifugation at 5000 r.p.m for 5 mins. Pelleted cells were then fixed by resuspending them gently in 1 mL of 3.7% (v/v) formaldehyde in 1× PBS prepared in nuclease-free water, followed by 30 min incubation at room temperature. Next, cells were centrifuged and washed twice in 1 mL of 1× PBS, followed by permeabilization in 1 mL of 70% ethanol (v/v) in nuclease-free water, with gentle mixing at room temperature for 1 hour. Afterward, they were washed once more in 1 mL of washing buffer (2× SSC solution in nuclease-free water containing 40% (w/v) formamide). Probing was done by treating the cells with hybridization buffer (consisting of 2× SSC, 40% (w/v) formamide, 10% (w/v) dextran sulfate, 2 mM ribonucleoside-vanadyl complex, 0.2 mg/mL BSA, and 1 mg/mL carrier *E. coli* tRNA, along with 1 μM of each fluorescent probe), overnight at 30 °C. Next, the cells were washed thrice with the washing buffer to remove excess probes, and subsequently resuspended in 2× SSC buffer. 3 μL of the cell suspension were sandwiched between the coverslip and 1% (w/v) agarose pads. In order to improve the photostability of Cy5 dye, the agarose pads were enriched with an oxygen scavenging system^29^ consisting of 2.5 mM protocatechuic acid (Sigma, prepared from a 100 mM stock stored frozen in water-NaOH, pH 8) and 0.05 U/mL protocatechuate 3,4-dioxygenase (OYC Americas).

### RT-qPCR measurements of mVenusNB expression

As in FISH experiments, overnight cultures were diluted 1:100 into fresh M9 medium and grown until mid-exponential phase. Next, cultures were treated with twice the volume of RNA-Protect bacteria reagent (Qiagen, Germany) at room temperature for 5 mins and centrifuged at 5000 r.p.m for 10 mins. An enzymatic lysis was performed on the pelleted cells with Lysozyme (10 mg/mL) in Tris-EDTA buffer (pH 8.0) and 10 % SDS. From these lysates, the total RNA content was extracted using the PureLink RNA Mini Kit (ThermoFisher) as per the kit manufacturer’s instructions. The RNA content and absorbance ratios A260/A280 nm and A260/A230 nm were quantified by a Nanodrop 2000 Spectrophotometer (Thermo Scientific). The ratio (2.0-2.1) indicated highly purified RNA.

DNA contamination was removed by treating the samples with DNase using the Turbo DNA-free kit (Invitrogen). Next, cDNA synthesis was performed from RNA through the High Capacity Reverse Transcription kit (ThermoFisher) as per the manufacturer’s instructions.

cDNA samples were mixed with qPCR master mix with the Power SYBR Green PCR mix (ThermoFisher) with primers (200nM) for the target and reference gene. The primer sequences for the target (mVenusNB) and reference (rrsA) genes were shown in supplementary table 2. Experiments were performed in a StepOne^TM^ Real-Time PCR system (ThermoFisher). The thermal cycling conditions were 50 °C for 10 minutes, 95 °C for 2 minutes, followed by 40 cycles of 95 °C for 15 seconds, 60 °C for 30 seconds, 72 °C for 40 seconds with the fluorescence being read after each cycle, and finally a melting curve from 90 °C through 70 °C at 0.3 °C intervals and 1 second dwell time. All samples were performed in 3 technical replicates, and for each condition, No-reverse-transcriptase and no-template controls were used to crosscheck non-specific signals and contamination. qPCR efficiencies of these reactions were greater than 95%. The fold change was calculated from the C_T_ values from target gene (normalized to the reference gene) and standard error, using Livak’s 2^-ΔΔCT^ method^30^.

### mRNA degradation rates

To measure the mVenusNB mRNA lifetimes in strains varying the operator sites (O_sym_, O_1_ and O_2_), we followed the procedure described in ref.^16^. From overnight cultures, cells were diluted 1:100 in a 20 mL volume of M9 succinate medium supplemented with 0.5X RPMI, grown at 37 °C with 200 r.p.m. Upon reaching OD_600_ ≅ 0.4, the culture was divided into two equal halves, with each half transferred to a new 50 mL tube and kept at 37 °C with aeration. Subsequently, 1.0 mL of culture was extracted from each tube and mixed with 3 mL of RNAprotect Bacteria reagent (Qiagen, Germany) to stabilize cellular RNA, serving as the t = 0 samples. Next, to inhibit transcription a final concentration of 500 μg/mL of Rifampicin was added to one of the culture tubes, while the culture without rifampicin served as a control. At two minute intervals after rifampicin addition (e.g., at t = 2, 4, 6, 8 minutes), 1.0 mL of culture was extracted from each tube and mixed with 3 mL of Qiagen RNAprotect Bacterial reagent. Subsequent total RNA extraction and quantitative real-time polymerase chain reaction (qRT-PCR) were carried out following the procedures outlined above. The relative mVenusNB mRNA levels in the rifampicin-treated sample were fitted to an exponential function, RNA = exp(-*γ*·t) where *γ* is the mRNA degradation rate and t is the time since rifampicin addition, yielding an estimate of *γ* = 0.79 min^-1^.

### Microscopy

Wide-field microscopy was employed to capture cells in both phase contrast and fluorescent channels. The optical setup consisted of a Nikon Ti2-E inverted microscope equipped with a 1.45/100x oil immersion objective lens (CFI Plan APO lambda, Nikon), a Spectra III light source from Lumencor for epi-fluorescence illumination, and a Kinetix camera from Teledyne Photometrix. Control of the setup was facilitated by micro-Manager 2.0 software^31^. For RNA measurements, we acquired Z-stacks of 9 Cy5 fluorescence images centered on the focal plane and separated by 200 nm, with a filter cube consisting of a FF660-Di02 excitation filter and a Semrock 692/40 emission filter. A single-phase contrast image was captured at the focal plane. All images were captured at an exposure time of 250 ms. For protein measurements, we used the same optical setup, and fluorescence images were captured using the mVenusNB channel with a filter cube consisting of an FF01-559/34 (Semrock) excitation filter, a T585lpxr (Chroma) dichroic mirror, and a T590LP (Chroma) emission filter. We acquired images at 200 ms exposure for both phase and fluorescence images.

### Image analysis

The image analysis pipeline consists of the following steps: cell segmentation, spot detection, and mRNA or mVenusNB quantification (see below for mVenusNB quantification). Cell segmentation was performed using the U-Net algorithm^32^. The segmentation was refined by imposing filters on cell area, width, and length; and each field of view was manually checked for mis-segmented cells. Fields of view with mis-segmented cells were excluded from further analysis.

For mRNA FISH experiments performed with *wt* LacI expression levels, spot detection proceeded similarly to previously published work^18^. Fluorescence images were subjected to a mild Gaussian filter (σ_XY_=1 pixel, σ_Z_=0.7 pixels), and local maxima were identified in 3D using Matlab’s imregionalmax function. Local maxima whose peak intensities fell under a threshold were discarded. This threshold was chosen such that at most a handful of spots were detected in the negative control sample (a negative control sample was run for each individual experiment). For each detected spot whose intensity passed the threshold, the spot intensity was determined by summing the pixels inside a circle of radius 5 pixels of the local maximum, in the XY plane in which the maximum was detected. To avoid double-counting, spots whose quantification radii overlapped in a particular XY plane were merged. The negative control pixel intensity was subtracted from all pixels while quantifying spot intensity.

To convert spot intensity to number of mRNA, the strain containing *wt lacI* and the *O*_*sym*_ operator was used. This strain is highly repressed, with on average 0.33 mRNA per cell, such that most detected spots are likely to contain a single mRNA. A histogram of detected spot intensities was created for the *O*_*sym*_ strain, and the single mRNA intensity was identified as the mode of the resulting intensity distribution (Supplementary Fig. 2).

With the single mRNA intensity in hand, the number of mRNA in each cell can be computed. The intensities from all detected spots in a cell are summed, and the resulting sum is divided by the single mRNA intensity, yielding the estimated number of mRNA in the cell. From the set of cells for each operator, the mean, Fano factor, and other statistics can be calculated. mRNA distributions in Fig. 2 and Fig. 3 were created by pooling the results of at least two independent experiments on separate days. Bootstrapped estimates of uncertainty in *k*_*a*_ and *k*_*d*_ (Fig. 3) were created as follows: for each operator, 1000 bootstrap samples were drawn from the pooled set of mRNA copy number observations. For each bootstrap sample, the *k*_*a*_ and *k*_*d*_ values were estimated as described above (“Calculation of *k*_*a*_ and *k*_*d*_ from mRNA statistics”). For error bars shown in Fig. 3b, the lower and upper error bar ranges represent the 10th and 90th percentile of bootstrapped *k*_*a*_ and *k*_*d*_ values, respectively.

For mRNA FISH experiments performed in Δ*lacI* strains, the summed fluorescence intensity of all pixels was calculated for each cell at the XY plane in which the cell was brightest, background fluorescence was subtracted using the negative control strain, and the background-subtracted summed fluorescence intensity was divided by the single mRNA intensity to yield the number of mRNA in each cell.

For mVenusNB expression experiments (Fig. 1c and d), the average mVenusNB channel pixel intensity was computed for each cell, and the average fluorescence of the negative control strain (without mVenusNB) was subtracted from this value. The background-subtracted mVenusNB intensity was averaged over all cells with a given operator to obtain a mean mVenusNB expression level for each operator. Error bars in Fig. 1d represent sem derived from pooling all cells with a given operator across experiments, drawing 10000 bootstrap samples from the pooled data set, and computing the standard deviation of the bootstrapped mean values.

### Stochastic simulations

Stochastic simulations in Fig. 1b were performed using code from Sanchez *et al*^33^. The steady-state probability distributions in Fig. 1b were computed from the chemical master equation by formulating the master equation in matrix form and numerically computing the eigenvectors of the matrix with eigenvalue equal to zero, also as described in Sanchez *et al*.

To generate Supplementary Fig. 8, investigating the effects of RNAP occlusion of the operator, this code was modified to make the association rate *k*_*a*_ dependent on the time since the most recent transcript production event. Specifically, the association rate was set to zero for a time *t*_*occlude*_ following each transcription event; after *t*_*occlude*_, *k*_*a*_ returned to its previously set value. Thus, *t*_*occlude*_ models occlusion of the operator during transcription initiation, during which time the operator is unavailable for binding. When constructing Supplementary Fig. 8, the association rate *k*_*a*,*sim*_ was modified for each operator such that the effective association rate when *t*_*occlude*_ = 1 s would approximately equal the experimentally measured *k*_*a*,*exp*_ for that operator. For instance, in the case of *O*_*sym*_, *r* = 24.6 min^-1^ (Fig. 2) and *k*_*a*,*exp*_ = 9.79 min^-1^. This means that, on average, 24.6 s of every minute are unavailable for LacI binding, so that the “true” association rate would be multiplied by a factor of (60-24.6)/60 = 0.59. Thus, to create the O_*sym*_ simulations in Supplementary Fig. 8, k_a,sim_ was set to *k*_*a*,*sim*_ = 9.79/0.59 min^-1^ = 16.6 min^-1^, which is indeed the inferred value for *k*_*a*_ when *t*_*occlude*_=0. An analogous procedure was used to determine *k*_*a*,*sim*_ values for *O*_*1*_ and O_2_ that, when taking into account an occlusion time *t*_*occlude*_ = 1 s, would yield inferred values close to the experimentally observed rates *k*_*a*,*exp*_ and *k*_*d*,*exp*_. For all three operators, this procedure worked well as inferred k_a_ and k_d_ values were close to the experimentally determined values when *t*_*occlude*_ = 1 s.

To generate Supplementary Fig. 9, a set of 50 simulated operators was created whose association rate per repressor 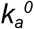 and dissociation rate *k*_*d*_ values were linearly spaced along the line defined in Marklund *et al* (Supplementary Fig. 9, green lines). In order to capture the effect of varying repressor copy number *n*_*repressor*_, the association rate *k*_*a*_ was assumed to be proportional to repressor copy number: 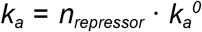. We examined 6 different repressor copy numbers, varying from 1 to 100 per cell. For a given operator and repressor copy number, the steady-state mRNA copy number distribution was computed numerically, again as described in Sanchez *et al*^33^, and a set of 1000 samples (i.e. comparable to performing mRNA FISH on a population of 1000 cells) was drawn from the distribution. The observed mRNA copy numbers in the simulated population were then analyzed according to the procedure in Fig. 2, yielding the “Inferred values” (blue dots). In order to estimate the effects of experimental noise, each mRNA was assigned a “fluorescence intensity” randomly chosen from a Gaussian distribution with mean 1 and standard deviation 0.4. The intensities from all mRNA within a particular cell were summed and rounded to the nearest integer, similarly to how experimental data is analyzed. This data was also analyzed according to the procedure in Fig. 2 (“Inferred values w exp noise”, orange dots). Basal transcription rate *r* = 22 min^-1^ and mRNA degradation rate γ = 0.80 min^-1^ in these simulations.

## Acknowledgments

We thank Johan Paulsson, Jinwen Yuan, Emil Marklund, and Daniel Camsund for their helpful discussions. We also thank I. Barkefors for critical reading of the manuscript. DJ acknowledges support from the Swedish Research Council (grant number 2020-05137). JE acknowledges funding from the ERC (advanced grant no. 885360), the Swedish Research Council (grant nos. 2016-06213 and 2018-03958), the Knut and Alice Wallenberg Foundation (grant nos. 2016.0077, 2017.0291, and 2019,0439.), and the eSSENCE e-science initiative. Computation and data management were enabled by resources provided by the Swedish National Infrastructure for Computing at UPPMAX, partially funded by the Swedish Research Council through grant agreement no. 2018-05973.

## Contributions

JE and DJ conceived and supervised the project. VK performed experiments. VK, SZ, and DJ analyzed data. DJ performed simulations. VK, JE, and DJ wrote the manuscript.

## Code and data availability

Code, experimental, and analysis data sets will be available at DOI:10.17044/scilifelab.26425576 upon manuscript acceptance.

