## Supplementary Figures for "Anti-correlation of LacI association and dissociation rates observed in living cells"

Supplementary Fig.1

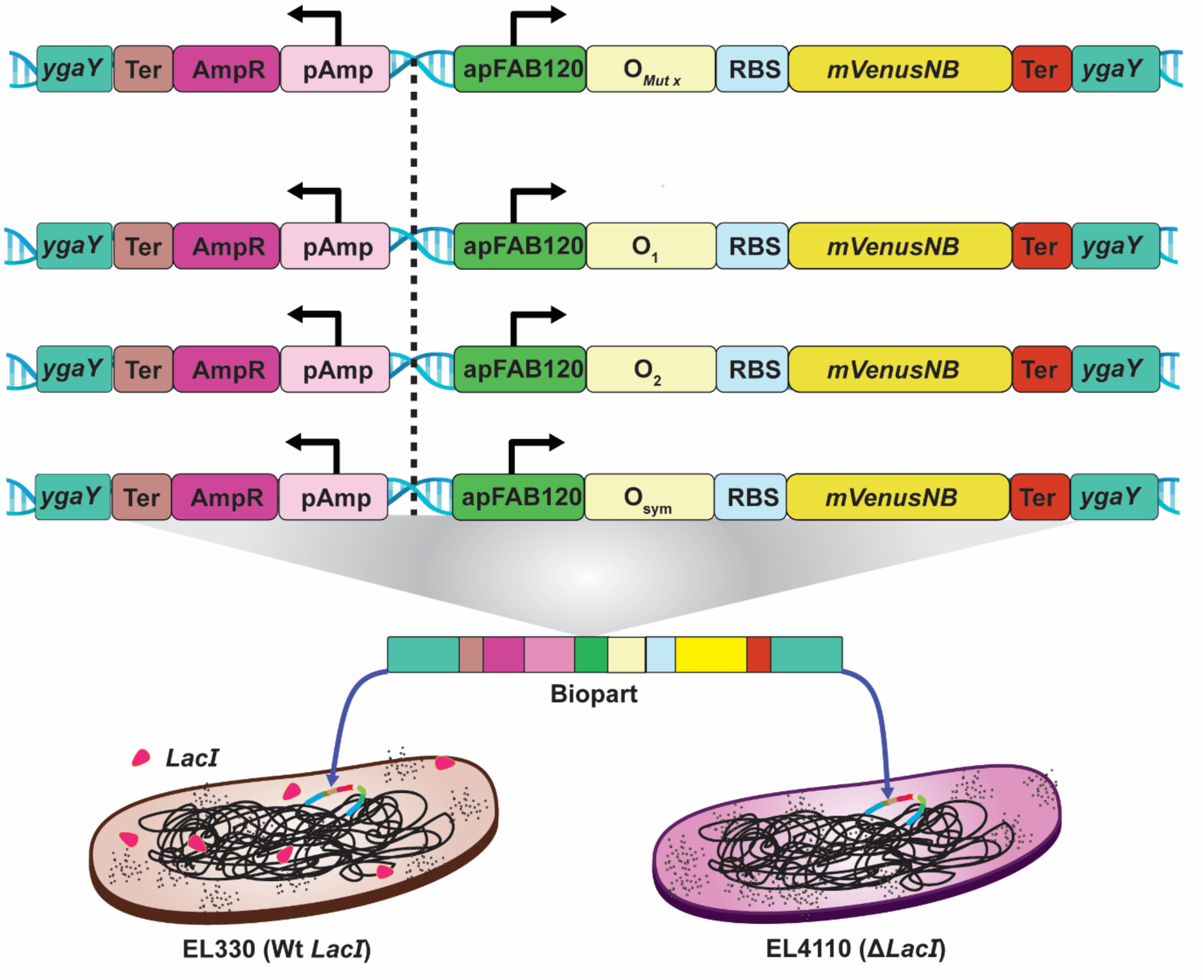


Supplementary Fig. 1: Schematic of promoter-operator Biopart and its insertion into the chromosome. The Biopart encodes the mVenusNB fluorescent protein expressed from the constitutive apFAB120 promoter. Downstream of the promoter, various LacI binding site ("Operator") mutants were introduced. Upstream of this promoter is a gene encoding ampicillin resistance, serving as a selection marker. This biopart is inserted into the *ygaY* locus in the chromosomes of two different host strains, which differ in the presence/absence of the *lacI* gene.

Supplementary Fig. 2

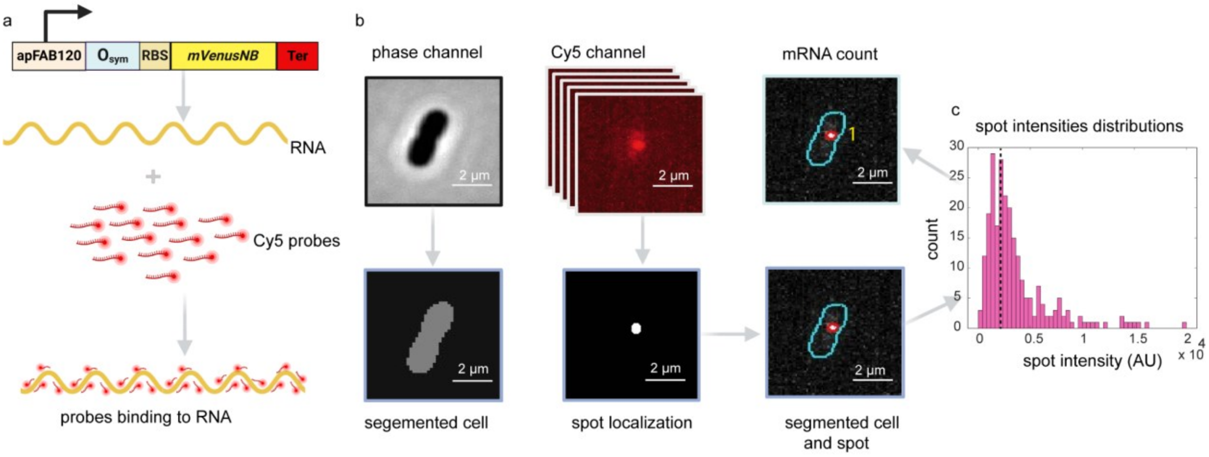


Supplementary Fig. 2: Quantification of single mRNA spot intensities. **a** Illustration of Biopart transcription and binding of Cy5 probes to mVenusNB mRNA. **b** Image acquisition of cells in the phase channel and fluorescence channel (top), cell segmentation, spot localization and segmentation from the Z stacks fluorescence images were performed using the custom-built pipeline in our lab (bottom). **c** Distributions of mRNA spot intensities from observed cells. The dotted line represents the inferred single mRNA spot intensity value (≈2400 AU).

Supplementary Fig. 3

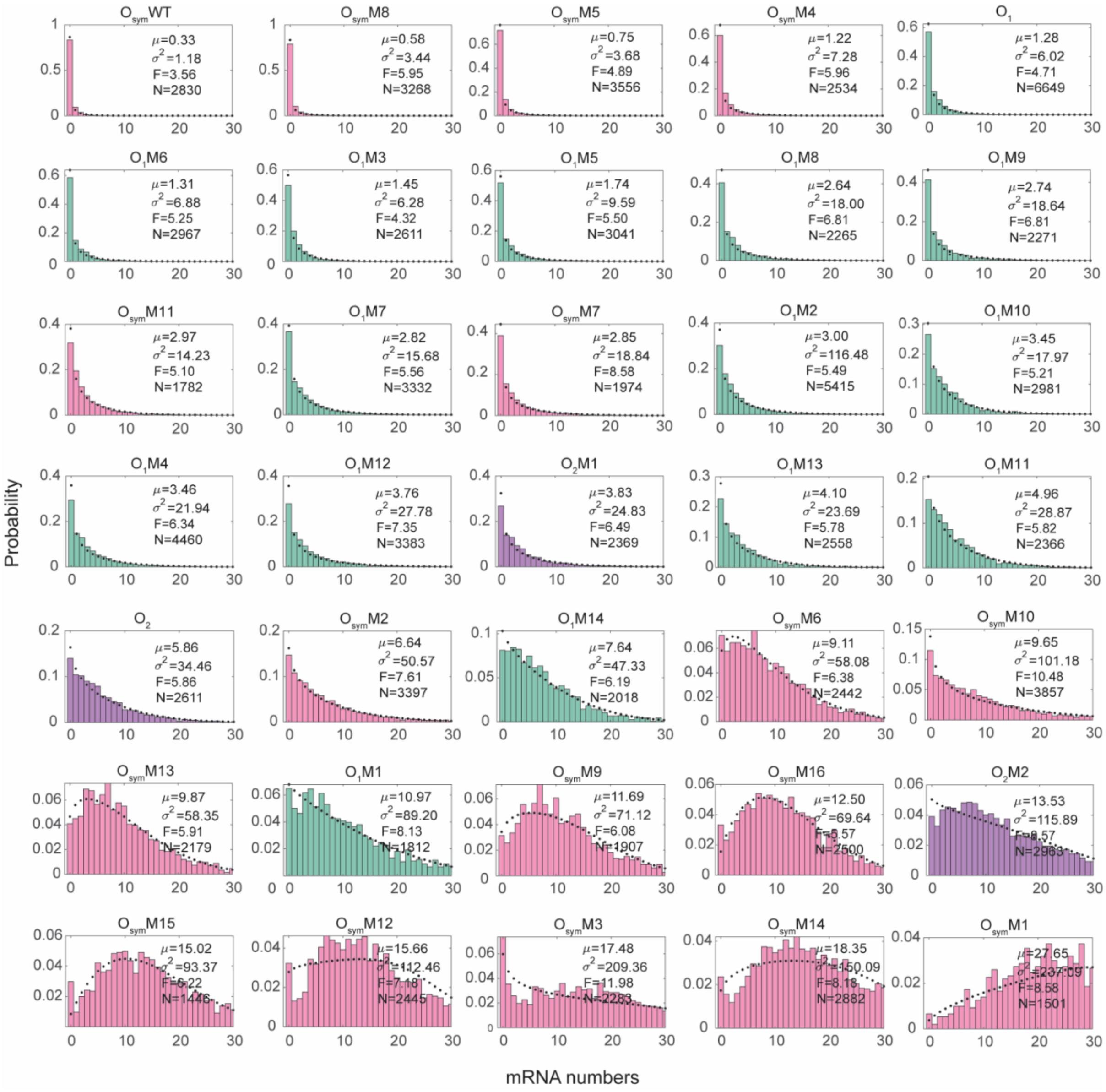

Supplementary Fig. 3: Observed mRNA copy number distributions in *wt lacI* strains, for all operators assayed in this work. For each operator, the probability mass function (pmf) corresponding to inferred $r$, $k_a$, and $k_d$ values is indicated with a dotted line.

Supplementary Fig.4

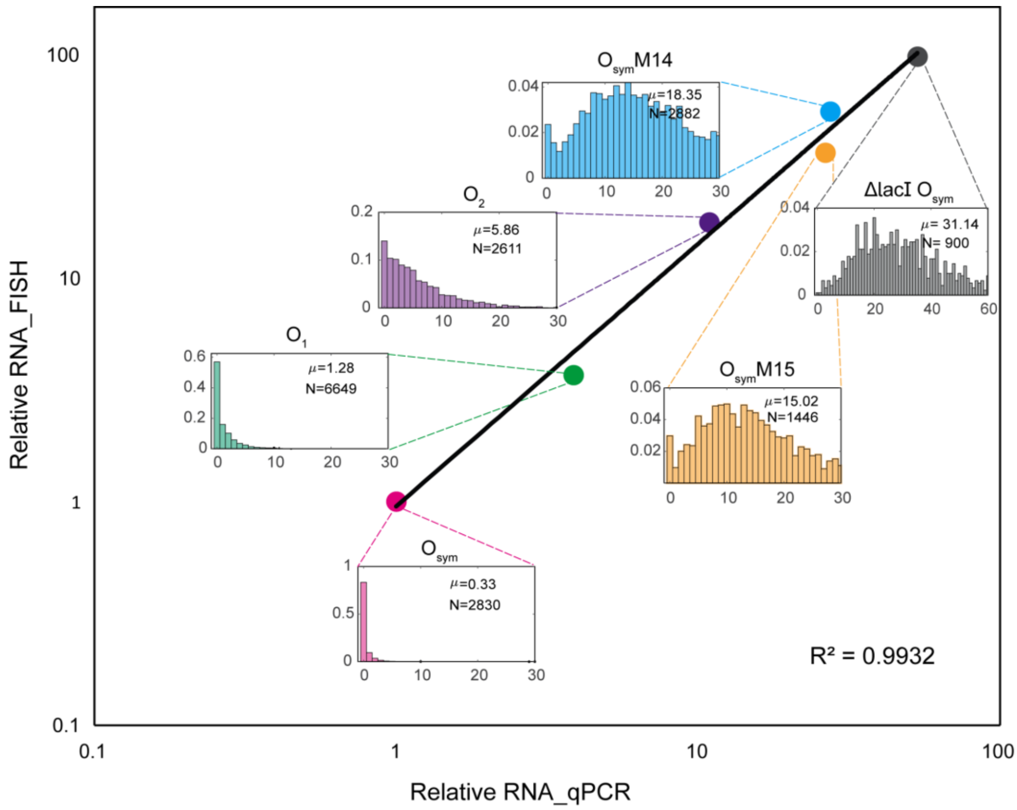


Supplementary Fig. 4: Comparison of mean mRNA FISH data with qPCR measurements. We selected six constructs ranging from low (0.33 mRNA/cell) to high (31.4 mRNA/cell) expression and compared mRNA FISH values with relative qPCR data. We find a positive correlation between these two independent measurements with an $R^2$ value of 0.9932.

Supplementary Fig. 5

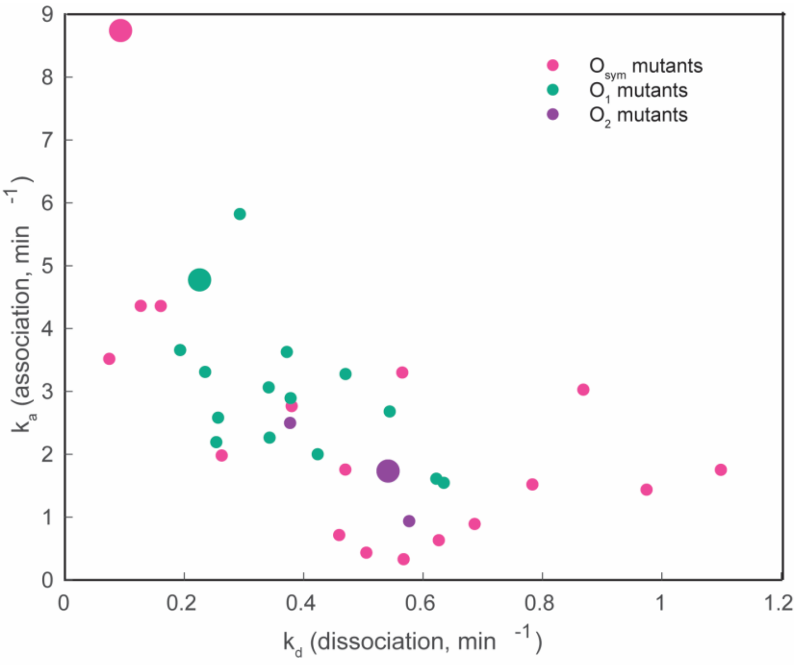


Supplementary Fig. 5: Inferred $k_a$ and $k_d$ values for all 35 operators without Fano factor correction.

Supplementary Fig. 6

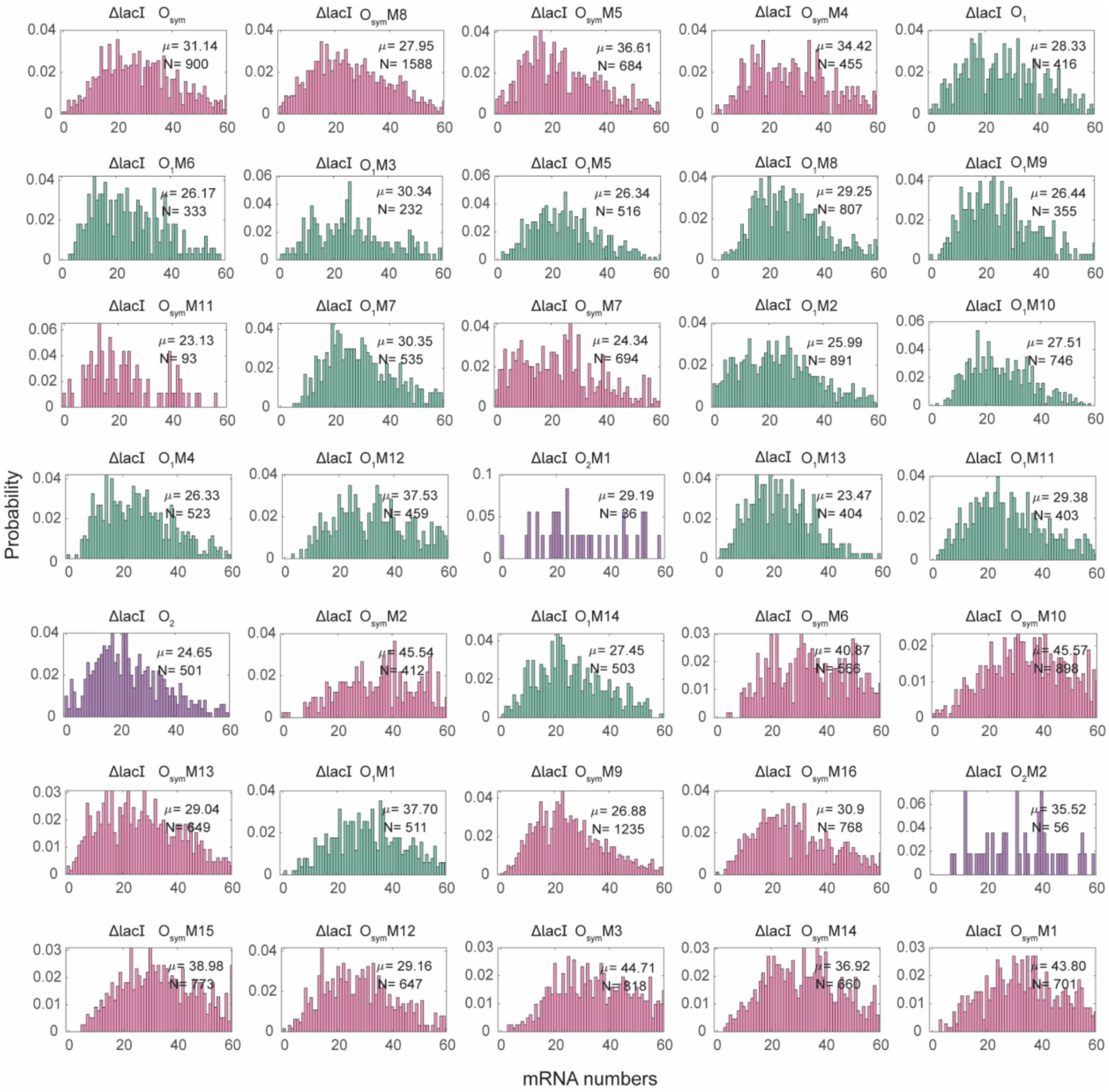


Supplementary Fig.6: Observed mRNA copy number distributions in *ΔlacI* strains, for all operators assayed in this work.

Supplementary Fig.7

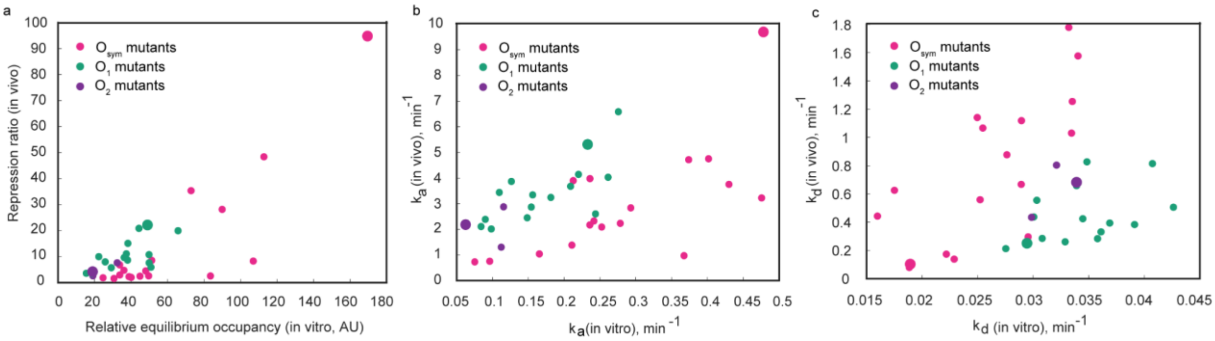


Supplementary Fig. 7: Comparison of in vivo and in vitro data. **a** The repression ratio (defined as the ratio of mRNA expression in the *ΔlacI* strain background to expression in the *wt lacI* strain background) is plotted against the relative *in vitro* occupancy measured in[1]. As expected, operators with higher occupancy display higher repression ratios. **b** and **c** The association rate $k_a$ and dissociation rate $k_d$ are compared in vivo and in vitro (again using data from[1].). The two measurements are correlated, but $O_{sym}$ mutants generally display slower association and faster dissociation than expected based on i*n vitro* values.

Supplementary Fig. 8

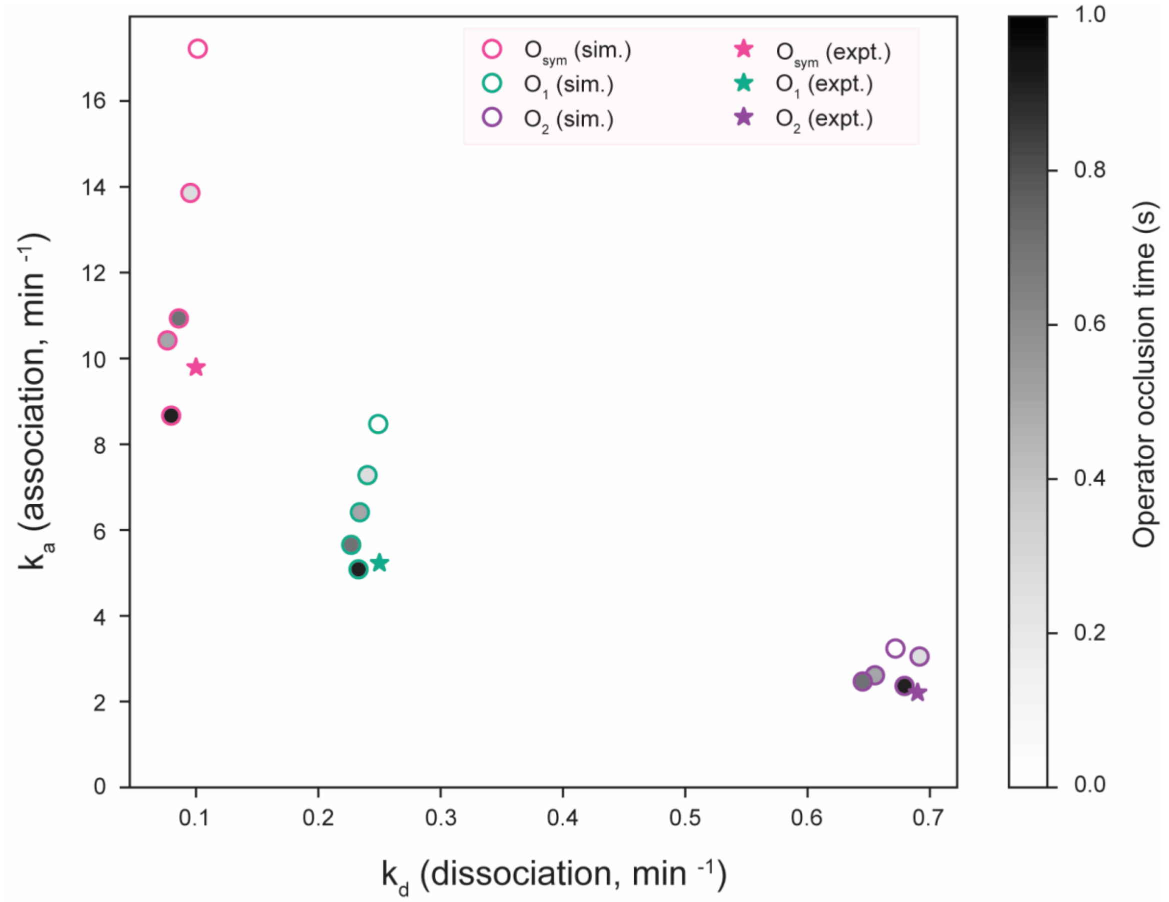


Supplementary Fig. 8: Effect of RNAP occlusion of LacI binding sites on observed association and dissociation rates. While clearing the promoter region, RNAP may temporarily inhibit LacI from binding. This “occlusion time” may in turn affect our calculations of LacI binding kinetics. To estimate the magnitude of this effect, we performed stochastic simulations as in Figure 1, with the modification that, upon production of a transcript, the association rate $k_a$ is set to zero for a time $t_{occlude}$ after the transcription event, in order to reflect the unavailability of the operator for LacI binding. Specifically, for each of the operators $O_{sym}$, $O_1$, and $O_2$, we simulated a population of 20 000 cells for 100 minutes, for occlusion times $t_{occlude}$ = 0, 0.25, 0.5, 0.75, and 1 seconds. At the end of the simulation, the mRNA copy number distribution over the 20 000 cells was computed, and $k_a$ and $k_d$ were estimated from simulated data using the procedure outlined in Figure 2 (circles; grayscale shading in circles reflects $t_{occlude}$ values). For each of the operators, the $k_a$ and $k_d$ values in the $t_{occlude}$=0 case were chosen such that estimated $k_a$ and $k_d$ values in the $t_{occlude}$=1 case would be close to the experimentally determined values (stars). Longer occlusion times lead to lower observed association rates, since LacI must bind in the intervals between transcript production events. Thus, it is possible that the experimentally-determined $k_a$ values reported in this work underestimate the "true" $k_a$ values, but this should not affect the observed anticorrelation between $k_a$ and $k_d$. In fact, these simulations indicate that the true spread in $k_a$ values may be larger than what we report.

### Supplementary Fig. 9

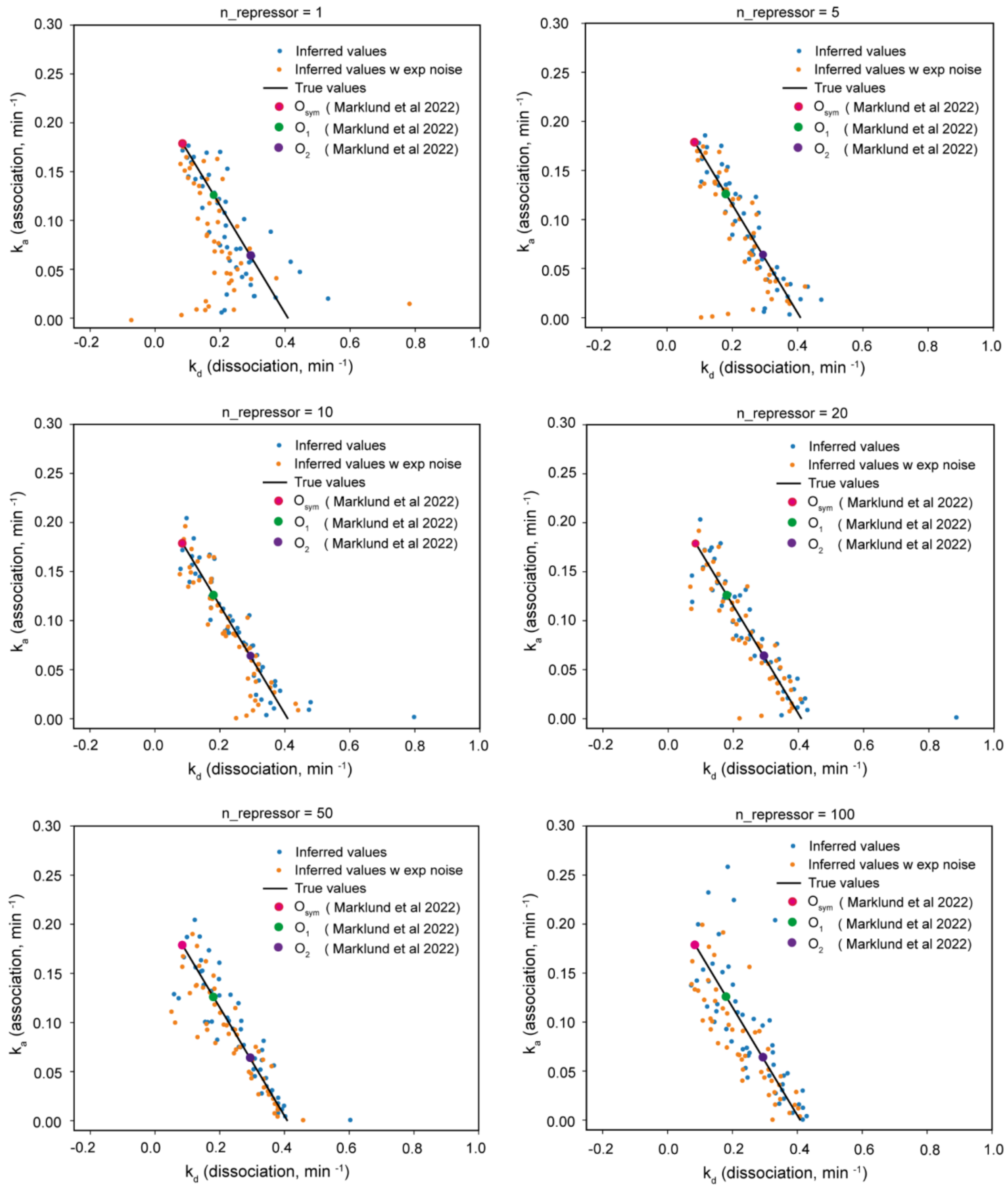


Supplementary Fig. 9: Analysis of simulated data for varying LacI concentrations. In order to assess the feasibility of the project, and to choose an appropriate LacI expression level, a set of 50 simulated operators was created whose association rate per repressor $k_a^0$ and dissociation rate $k_d$ values were linearly spaced along the line defined in Marklund *et al*[1] (solid black lines, "True values"). In order to capture the effect of varying repressor copy number $n_{repressor}$, the association rate $k_a$ was assumed to be proportional to repressor copy number: $k_a = n_{repressor} \cdot k_a^0$. We examined 6 different repressor copy numbers, varying from 1 to 100 per cell. For a given operator and repressor copy number, the steady-state mRNA copy number distribution was computed numerically, again as described in Sanchez *et al*[2], and a set of 1000 samples (i.e. analogous to performing mRNA FISH on a population of 1000 cells) was drawn from the distribution. The observed mRNA copy numbers in the simulated population were then analyzed according to the procedure in Fig. 2, yielding the "Inferred values" (blue dots). To facilitate comparison between different simulated repressor copy numbers, the association rate per repressor $k_a^0$ is plotted on the y axis rather than association rate $k_a$. Note that $k_a$ and $k_d$ values in this figure are based on *in vitro* data and therefore do not perfectly match *in vivo* estimates from Figs. 2 and 3, although trends are the same. In order to estimate the effects of experimental noise, each mRNA within each simulated cell was assigned a "fluorescence intensity" randomly chosen from a Gaussian distribution with a mean 1 and a standard deviation 0.4. The intensities from all mRNA within a particular cell were summed and rounded to the nearest integer, similarly to how experimental data is analyzed. This data was also analyzed according to the procedure in Fig. 2 ("Inferred values w exp noise", orange dots), and evidently, this level of variability in the single mRNA intensity did not lead to notably worse inference of LacI kinetics. Inference is most accurate at circa 10 repressors per cell, similar to the wild-type LacI expression[3]. For lower or higher LacI expression, the inferred values are less tightly clustered around the true values. Basal transcription rate $r$ = 22 $min^{-1}$ and mRNA degradation rate $\gamma$ = 0.8 $min^{-1}$ in these simulations.
